# Characterization of *NvLWamide-like* neurons reveals stereotypy in *Nematostella* nerve net development

**DOI:** 10.1101/145623

**Authors:** Jamie A Havrilak, Dylan Faltine-Gonzalez, Yiling Wen, Daniella Fodera, Ayanna C. Simpson, Craig R. Magie, Michael J Layden

**Affiliations:** Lehigh University Department of Biological Sciences; Quinnipiac University Department of Biological Sciences

**Author notes:** Corresponding Author –, Lehigh University Department of Biological Sciences, 111 Research Drive, Iacocca Hall Rm B-217, Bethlehem, PA 18015.

**Keywords:** *Nematostella*, Transgenenic, Neurogenesis, Nerve net, LWamide

## Abstract

Cnidarian nervous systems are traditionally described as diffuse nerve nets lacking true organization. However, there are examples of stereotypical structure in the nerve nets of multiple cnidarian species that suggest nerve nets are organized. We previously demonstrated that the *NvashA* target gene *NvLWamide-like* is expressed in a small subset of the *Nematostella* nerve net and speculated that observing a few neurons within the developing nerve net would provide a better indication of potential stereotypy. Here we document *NvLWamide-like* expression more systematically. *NvLWamide-like* is initially expressed in the typical neurogenic salt and pepper pattern within the ectoderm at the gastrula stage, and expression expands to include endodermal salt and pepper expression at the planula larval stage. Expression persists in both ectoderm and endoderm in adults. We generated an *NvLWamide-like::mCherry* transgenic reporter line to visualize the neural architecture. *NvLWamide-like* is expressed in six neural subtypes identifiable by neural morphology and location. Upon completing development the numbers of neurons in each neural subtype are minimally variable between animals and the projection patterns of each subtype are consistent. Between the juvenile polyp and adult stages the number of neurons for each subtype increases. We conclude that cnidarian nerve nets are organized, develop in a stereotyped fashion, and that one aspect of generating the adult nervous system is to modify the juvenile nervous system by increasing neural number proportionally with size.

## Introduction

Possessing a centralized nervous system (CNS) is a unifying feature of bilaterians (chordates, insects, nematodes, acoels, annelids, mollusks, echinoderms, etc.). Details regarding the origin of bilaterian central nervous systems are still under debate, but most believe an ancestral nerve net preceded the organized condensation of neurons typical of a bilaterian CNS (Arendt et al., 2008; Ghysen, 2003; Hartenstein and Stollewerk, 2015; Watanabe et al., 2009); Galliot et al., 2009). Cnidarians (e.g. jellyfish, corals, anemones, *Hydra,* etc.) represent the sister taxon to the bilaterians, and their nervous systems are comprised of a nerve net wherein neurites emanating from neural soma scattered throughout the body form a “net” encompassing the organism (Dunn et al., 2008; Hejnol et al., 2009; Rentzsch et al., 2016a). The phylogenetic position of cnidarians in relationship to bilaterians together with the fact that they have a nerve net, puts this group in a unique position to inform and potentially test hypotheses about the origin and evolution bilaterian centralized nervous systems. However, very little is known about the developmental patterning of cnidarian nerve nets, which makes it difficult to compare components of cnidarian nerve nets to their putative bilaterian counterparts. In particular, it isnt clear if stereotypical development of cnidarian nerve nets occurs. Thus, determining the stereotypy of nerve net development and identifying distinct neural subtype markers is important to then assess how nerve nets are patterned.

Cnidarian nerve nets have traditionally been described as being diffuse and disorganized in the sense that there is minimal reproducible architecture that is obvious from animal to animal within a given species. However, a number of observations suggest cnidarian nerve nets are more organized than usually appreciated. First, there are obvious structures identified in many cnidarian nervous systems. For example, the sea anemone *Nematostella vectensis* has clearly visible longitudinal tracts that run the length of the oral-aboral axis and are stereotypically positioned over each mesentery structure (Layden et al., 2016b; Marlow et al., 2009; Nakanishi et al., 2012; Rentzsch et al., 2016a). In *Hydra* the foot of the animal has been identified as an organizing center for distinct behaviors, which indicates unique functions are regionally assigned within the nerve net. There are distinct subsets of neurons with particular spatial locations that are identifiable in *Hydra* and *Nematostella* (Anderson et al., 2004; Grimmelikhuijzen and Spencer, 1984; Koizumi et al., 2004; Marlow et al., 2009). A nerve ring containing at least 4 different neuronal subsets has been identified at the base of tentacles in *Hydra oligactis* and thought to be involved in feeding behaviors (Koizumi et al., 2015). Two marginal nerve rings have been identified in the jellyfish *Aglantha* (Mackie, 2004; Donaldson et al., 1980; Roberts and Mackie, 1980). Hydrozoan nerve rings are visualized with various different antibodies that do not overlap/colocalize, suggesting that distinct neuronal subpopulations populate the nerve ring, although the number of neurons within and organization of these subpopulations is not clear (Koizumi et al., 2015). Altogether, these observations across many cnidarian species hint at more patterning and organization within a typical cnidarian nerve net than once thought.

The sea anemone *Nematostella vectensis* is an anthozoan cnidarian model system that has grown in popularity due to its ease of culture and the ever-growing repertoire of genomic tools available (Darling et al., 2005; Layden et al., 2016b; Putnam et al., 2007). While Cnidarians posses both ectodermal and endodermal nerve nets, the developmental origin varies between species. In *Nematostella,* neurogenesis occurs in both the ectoderm and endoderm (Nakanishi et al., 2012). Studies primarily conducted in *Nematostella* suggest that cnidarian and bilaterian neural patterning likely share a common evolutionary origin. (Layden et al., 2016b; Rentzsch et al., 2016b). For instance, MEK/MAPK (Layden et al., 2016a), Wnt (Leclère et al., 2016; Marlow et al., 2013; Sinigaglia et al., 2015), BMP (Saina et al., 2009; Watanabe et al., 2014) and Notch (Layden and Martindale, 2014; Richards and Rentzsch, 2015), all key regulators of bilaterian neurogenesis, have similar roles in *Nematostella.* Additionally, key neurogenic transcription factors such as *NvsoxB(2)* and *Nvath-like* are expressed in proliferating neural progenitor cells (Richards and Rentzsch, 2015; 2014). The bHLH proneural transcription factor *NvashA* is necessary and sufficient to promote neurogenesis including the formation of individual neuronal subtypes (Layden et al., 2012). It has been suggested that the conserved neurogenic programs described above coordinate with axial patterning to generate distinct neural subtypes, similar to mechanisms that pattern bilaterian central nervous systems (Layden et al., 2012; Leclère et al., 2016; Rentzsch et al., 2008; Watanabe et al., 2014). However, an understanding of cnidarian neural architecture or the patterning of distinct neural subtypes in these organisms is lacking, and thus with few exceptions there are not clear neural subtypes that can be used to screen for neural patterning defects.

Our previous efforts identified the *NvLWamide-like::mcherry* transgenic line (Layden et al., 2016a). We speculated that the neural number and neurite projection patterns were consistent from animal to animal, which if true would suggest development of nerve nets is at least in some cases stereotyped. Here we characterize the *NvLWamide::mcherry* transgenic line in more detail in order to better determine the development and organization of the juvenile and adult nervous system. We identified at least six neuronal subtypes described by *NvLWamide::mcherry* that are consistently found at particular locations in the polyp and with distinct neurite morphologies. By the juvenile polyp stage the number of each neural class is predictable from animal to animal and these subtypes persist through adult stages. Interestingly, the number of neurons for each individual neuronal subtype increases with age, and can in some cases be easily quantified revealing a positive correlation between neural number and body length. These combined data imply that the *Nematostella* nervous system develops and is modified in a stereotypical fashion, which contradicts the notion that nerve nets loosely organized structures.

## Materials and Methods

### Animal care

All animals were maintained in 1/3X artificial seawater (ASW) with a pH of 8.1–8.2, with weekly water changes. Embryos were either grown at 25°C, 22°C or 17°C to the desired stages, and juvenile polyps were maintained at room temperature in the dark. Adult *Nematostella* were housed in the dark at 17°C, were fed brine shrimp 4 times a week, and were given pieces of oyster 1 week prior to spawning. Spawning was induced by changing the light cycle (Fritzenwanker and Technau, 2002; Hand and Uhlinger, 1992). Generation of *NvLwamide-like* transgenic animals was described by Layden et al, 2016 (Layden et al., 2016a). All transgenic animals were from a stable F1 line, and since there was no visible difference in mCherry expression in heterozygous versus homozygous animals both were utilized in these studies.

### In situ hybridization and immunohistochemistry

Protocols for fixation, *in situ* probe synthesis, and both immunohistochemical or double fluorescent *in situ* hybridization were performed as previously described (Wolenski et al., 2013). The *NvLWamide-like* probe was previously described (Layden et al., 2012). The mCherry probe was amplified using the following primers: mCherry Forward: gcaagggcgaggaggacaac; mCherry Reverse: cttgtacagctcgtccatgccg. Cloning and probe synthesis were carried out as previously described (Wolenski et al., 2013).

Immuno-staining was performed as previously described ((Nakanishi et al., 2012; Wolenski et al., 2013)) with the following modifications. Incubations in primary and secondary antibodies were done overnight at 4°C. Anti-DsRed (Clontech Living Colors DsRed Polyclonal Antibody, 632496, Antibody registry ID: AB_10013483) was diluted 1:100 in blocking reagent (4% goat serum in PBS + 0.2% Triton X-100 +1% BSA). Secondary Alexa Fluor 555 antibody (Thermo Fisher Scientific, A21428) was used at 1:500 in 1x blocking reagent.

### Quantification of *mCherry*/dsred and *NvLWamide-like* positive cells

To quantify the number of *mCherry/α-dsred* positive cells that colabel with *NvLWamide-like* positive cells the embryos were mounted to capture z-stacks of lateral views with a 1uM step size using a Zeiss LSM 880 (Carl Ziess) confocal microscope. Cells were then scored as single or double positive using the Imaris (Bitplane) imaging software.

### Quantification of NvLWanide-like::mcherry-expressing neurons/ neuronal subtypes

NvLWamide-like::mCherry expressing juvenile polyps were relaxed in 7.14% (wt/vol) MgCl2 in 1/3 X ASW for 10 minutes at room temperature. Animals were either examined and quantified live, subjected to live imaging, or imaged following a light fixation and phalloidin stain. Animals were fixed in 4% paraformaldehyde and 0.3% glutaraldehyde (in 1/3 X ASW) for 1 minute, followed by fixation in 4% paraformaldehyde (in 1/3 X ASW) for 20 minutes at room temperature on a rocker. Animals were then washed in PBS + 0.2% Triton X-100 for 5x5 minute washes, then stained with Alexa Fluor 488 phalloidin (Life Technologies, A12379, 3:200) in PBS + 0.2% Triton X-100 for 1 hour. Animals were washed for 5x5 minutes and mounted in 90% glycerol for immediate imaging.

### Imaging

DIC images of *in situ* hybridization, and live images of NvLWamide-like::mCherry juvenile polyps and adult *Nematostella* were taken on a Nikon NTi with a Nikon DS-Ri2 color camera and the Elements software (Nikon). Confocal images of live and fixed/stained *Nematostella NvLWamide-like::mCherry* embryos and juvenile polyps were obtained using a Zeiss LSM 880 with LSM Zen software (Carl Ziess) or a Nikon C1 with EZ-C1 software (Nikon). Confocal images were processed using Imaris 8.2 software (Bitplane LLC) and 3D images were constructed from a composite of serial optical sections (z-stacks). Final images were cropped using Illustrator and/or Photoshop (Adobe Inc.).

## Results

### NvLWamide-like expression is scattered throughout the ectoderm and endoderm during development

*NvLWamide-like* is expressed in a subset of the nervous system, but its detailed spatiotemporal distribution has not been described (Layden et al., 2012). We used mRNA *in situ* hybridization to better characterize expression of *NvLWamide-like* during development (Figure 1). *NvLWamide-like* mRNA was first detected in a scattered pattern in the aboral hemisphere of the embryonic ectoderm at the early gastrula stage (Figure 1A), which recapitulated the previously published gastrula expression pattern (Layden et al., 2012). The ectodermal expression is detected throughout development (Figure 1B–E, black arrowheads), but the distribution of cells is dynamic. For example, there was a notable decrease in the number of *NvLWamide-like* expressing cells in the aboral ectoderm between the late gastrula and planula stages (compare Figures 1A and 1B). Also during planula development ectodermal expression extends orally and can be detected in the developing pharynx (Figure 1B and C, white arrowheads). By juvenile polyp stage, ectodermal *NvLWamide-like* expression is detected throughout the entire ectodermal body column and in the tentacles (Figure 1D–E).

**Figure 1.**
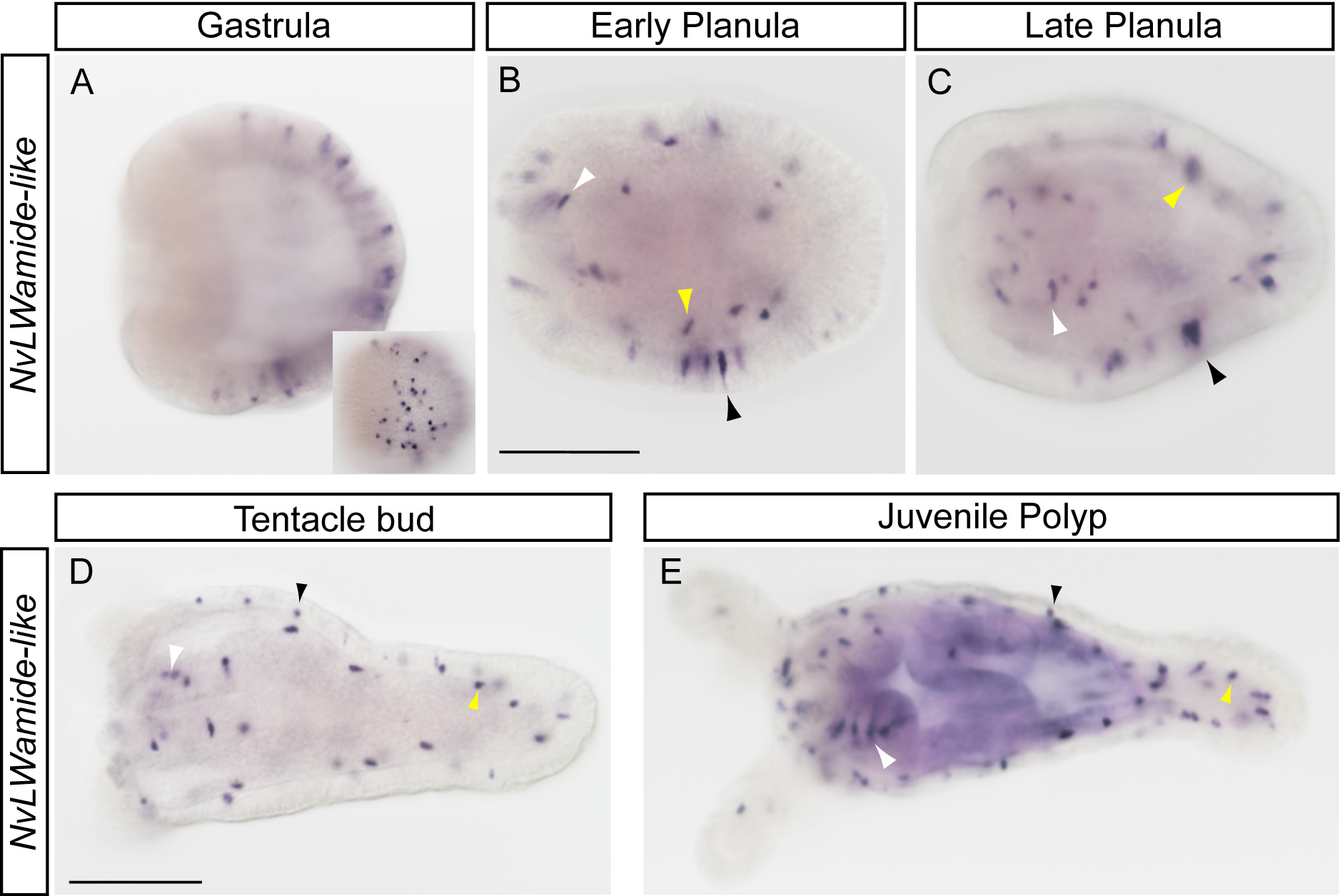
Expression of *NvLWamide-like* during development. **(A)** *NvLWamide-like* mRNA is first detected in ectodermal cells during the gastrula stage in a salt-and-pepper pattern. **(B-E)** Salt-and-pepper ectodermal *NvLWamide-like* expression is maintained in the early and late planula (B and C, black arrowheads), tentacle bud polyp (D, black arrowhead) and juvenile polyp (E, black arrowhead). *NvLWamide-like* expression is detected in the presumptive endoderm by the early planula stage (B, yellow arrowhead) and expression is maintained throughout development (C-E, yellow arrowheads). *NvLWamide-like* positive cells are observed in the pharyngeal ectoderm in the early planula stage (B, white arrowhead) and continue to be detected in the pharyngeal region throughout development (C-E, white arrowheads). *NvLWamide-like* cells are also present in the tentacles of juvenile polyps (E). All images are lateral views with the oral end towards the left. Inset is a surface view of the gastrula embryo in A. Scale bars = 100μm.

*NvLWamide-like* is also expressed in small number of endodermal cells. Endodermal expression is first detected in early planula stage animals (Figure 1C, yellow arrowhead). Endodermal expression is maintained throughout development (Figure 1C–E, yellow arrowheads) and into adult stages (see below). Additionally, endodermal cells are associated with the mesenteries and longitudinal tracks that run the length of the oral-aboral axis (Figure 2D) (Layden et al., 2016b; 2016a; Nakanishi et al., 2012).

**Figure 2.**
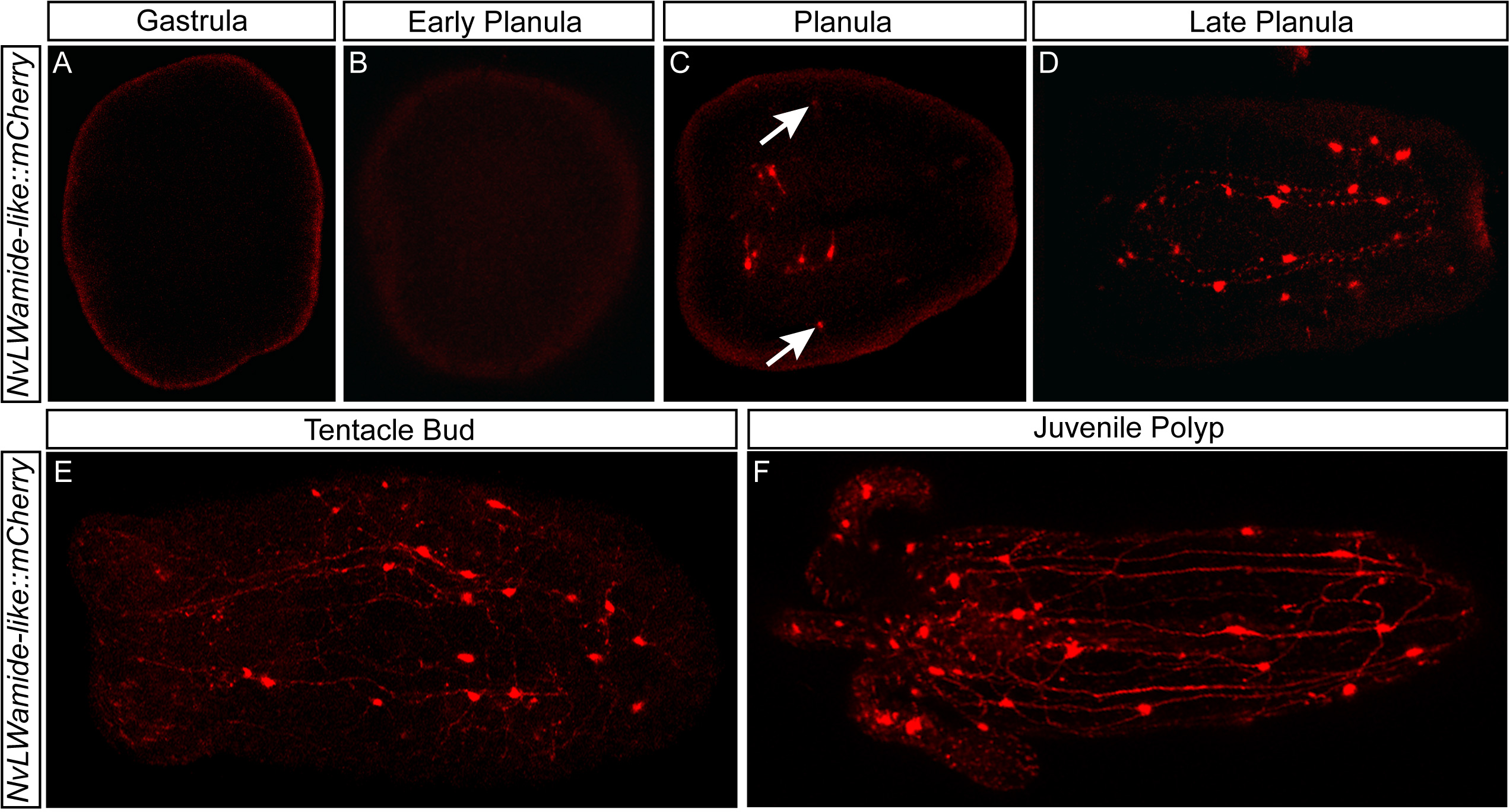
Visualization of NvLWamide-like::mCherry-expressing neurons during development. **(A-B)** *NvLWamide-like::mCherry* expression is not visible during the gastrula (A) or early planula (B) stages of development. **(C)** *NvLWamide-like::mCherry* positive neurons are first observed in the mid-planula stage, when neurons around the pharynx and within the ectoderm (arrows) become visible. **(D)** By the late planula stage expression can be seen in the longitudinal neurons (note neurites extending along the oral-aboral axis). **(E)** By the tentacle bud stage, more numerous lateral connections can be seen between longitudinal neurites. **(F)** The number of NvLWamide-like::mCherry positive neurons continues to increase into the juvenile polyp stage. Images depict *in vivo* mCherry expression in live animals.

### NvLWamide-like::mCherry expression recapitulates endogenous NvLWamide-like mRNA expression during polyp stages

To better determine what cell types express *NvLWamide,* we generated a stable F1 *NvLWamide::mCherry* transgenic reporter line, and either live imaged animals, or imaged animals that were lightly fixed and co-stained with phalloidin (see materials and methods).

We found that mCherry expression followed the expression dynamics of the endogenous *NvLWamide* transcript (compare Figures 1 and 2) with the exception that mCherry positive cells are not readily detected in gastrula and early planula stage animals (Figure 2A and B). *NvLWamide-like::mCherry* neurons are first detected in the ectoderm and in the pharynx of the planula (Figure 2C). Tentacular and endodermal expression of mCherry is detected at similar stages to those observed for the endogenous transcript (Figure 2D–F).

Because of the lack of mCherry transgene expression during early development we wanted to confirm that the *NvLWamide-like::mcherry* transgene faithfully recapitulates *NvLWamide-like* expression. To determine the degree of overlap between transgenic and endogenous *NvLWamide-like* expression we performed fluorescent *in situ* hybridization (FISH) and α-dsRed immunofluorescence to label endogenous *NvLWamide-like* and mCherry protein respectively. We also performed double FISH to compare endogenous *NvLWamide-like* to *mcherry* transcript co-expression. At gastrula and early planula stages no α-dsRed staining was detected (Figure 3A and B), consistent with our observation that mCherry is not detected at these stages. α-dsRed was first detected by immunofluorescence in the planula stage and we observed a range from 75–100% of α-dsRed positive cells co-expressing *NvLWamide-like* (N= 5animals, average co-expression = 84.3%) (Figure 3C), but many *NvLWamide-like* positive cells are still negative for α-dsRed. By polyp stages we observed a range of 80–91% of α-dsRed positive cells co-expressing *NvLWamide-like* (N=7 animals) (Figure 3D). We speculated that the observed delay in mCherry co-expression during early stages reflected a delay in mCherry protein maturation rather than a lack of co-expression at early stages. To address this hypothesis, *NvLWamide::mCherry* animals were fixed and double fluorescent *in situ* hybridizations were performed to label both the endogenous *NvLWamide-like* and the *mcherry* transcript. We were unable to detect *mcherry* in gastrula, but did detect *mcherry* in the early planula stages, and observed that 100% of *mcherry* positive cells co-expressed *NvLWamide* (Figure 3E–G)(N=6 animals). Similarly 100% of *mcherry* positive cells at polyp stages also showed 100% co-expression with *NvLWamide-like* (Figure 3H–J). Our data suggest that there is a delay in transgene expression compared to the endogenous *NvLWamide-like* expression, and that some of the gastrula *NvLWamide-like* positive cells do not express the transgene. However, by polyp stages there is effectively a 1:1 correlation of *NvLWamide-like* and *mcherry* expression indicating that our transgene faithfully recapitulates endogenous expression in the mature nervous system.

**Figure 3.**
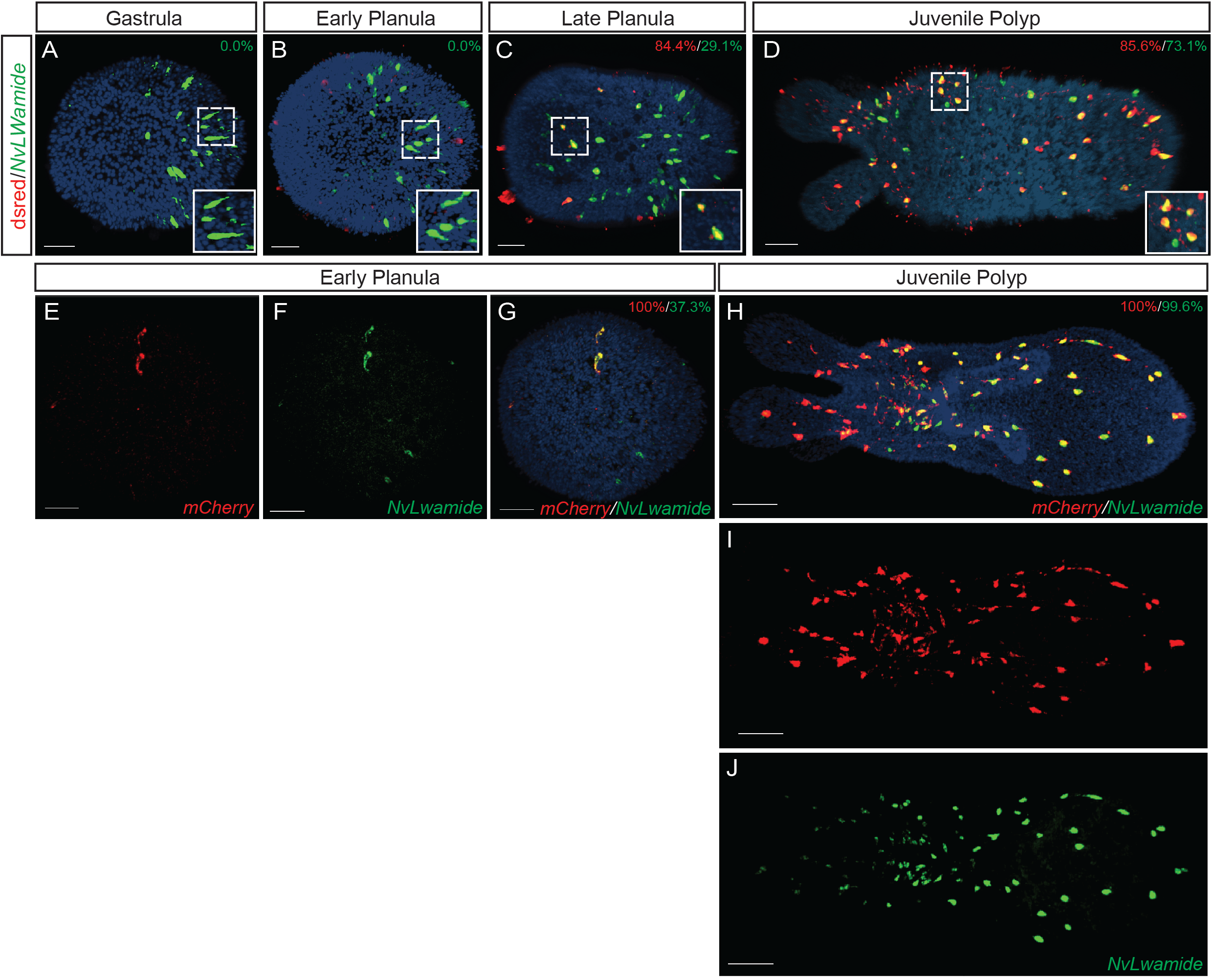
Colocalization of mCherry transgene with endogenous NvLWamide. **(A, B)** The mCherry transgenic protein, labeled with a-dsred, is not detected during gastrula and early planula stages. **(C)** By late planula stages 84.3% of the a-dsred positive cells are also *NvLWamide* positive (red) while only 29.1% of the *NvLWamide* mRNA positive cells are also a-dsred positive (green) **(D)** At the juvenile polyp stage 85.6% of the a-dsred positive cells are also *NvLWamide* mRNA positive (red) while 73.1% of the *NvLWamide* mRNA positive cells colabel with **a**-dsred (green). **(E-G)** Analysis of the *mCherry* transgenic mRNA showed that 100% of the mCherry positive cells are also *NvLWamide* positive at the gastrula stage and **(H-J)** 100% colocalization of the two at the juvenile polyp stage. Percentages in the top right corners indicate the percentage of *a-dsred/mCherry* positive cells which are also NvLWamide positive (red) and the percentage of NvLWamide mRNA positive cells which are also *a-dsred/mCherry* positive (green). For *NvLWamide* mRNA and a-dsred colabeling experiments n=5, 6, 5, 7 for early gastrula, gastrula, planula, and juvenile polyp respectively. For *NvLWamide* and *mCherry* mRNA colabeling experiments n=6 for gastrula and n=5 for juvenile polyps. All images are lateral views with the oral end towards the left. Scale bars = 30μm.

### NvLWamide-like::mCherry positive neurons represent distinct neuronal subtypes that develop in a stereotypic manner

We next characterized the neuronal subtypes that express *NvLWamide-like* during development by analyzing the distribution of and projection patterns present in different NvLWamide-like::mCherry-expressing cells. Initially, 12 day old juvenile polyps were analyzed to determine if the pattern of *NvLWamide-like* neurons were consistent between animals at the end of embryonic and larval development. We were able to identify six neural subtypes that we have assigned the identity of *NvLWamide-like* pharyngeal, -mesentery, -longitudinal, -tentacular, -tripolar, and -quadripolar neurons (Figures 4–6; Supplementary Figures 1 and 2; Supplemental movie 1). We also identified mCherry positive nematosomes, which are easiest to observe embedded in the egg mass during spawning (Figure 6K and L). Eighteen day old (Supp. Fig. 2A) and twenty three day old (Supp. Fig. 2B) polyps showed no changes in neural number and no additional neural subtypes compared to twelve day old polyps (Compare Supplemental Figure 2A and B with Figures 4A and 7A). However, by adult stages the same neural subtypes identified juvenile polyps were present but were increased in number (Figure 7). Below we describe each neuronal class in detail. We conclude that the neurons present in the juvenile polyp represent the complement of neurons specified during embryonic and larval development and that additional neurons are not born until after the polyp begins to grow once it starts to feed.

**Figure 4.**
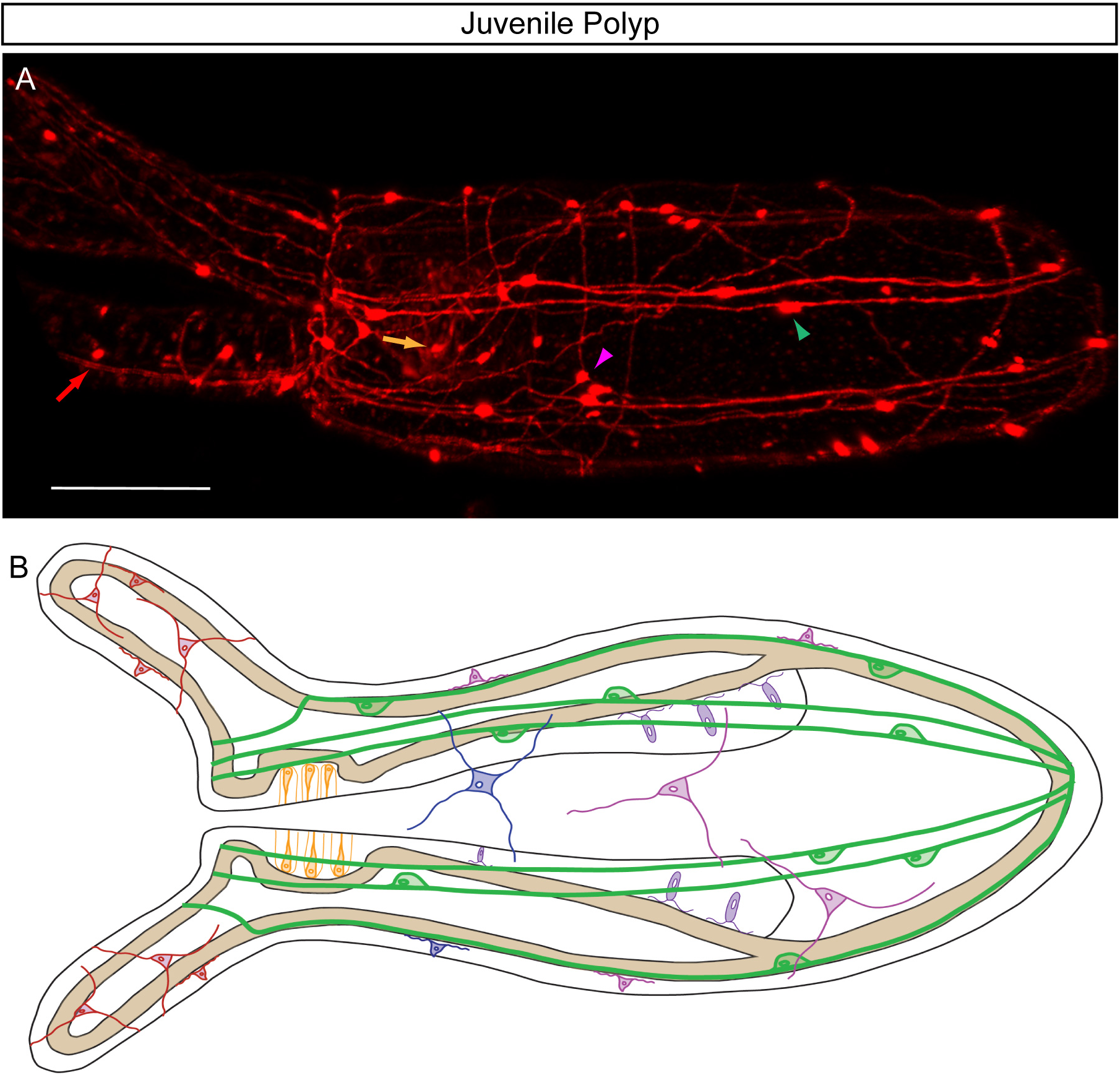
Overview of NvLWamide-like::mCherry positive neurons. **(A)** 3D projection of live confocal images of the NvLWamide-like::mCherry transgenic line identifies neurons in the juvenile polyp *in vivo.* Tentacular (red arrow), pharyngeal (orange arrow), tripole (magenta arrowhead), and longitudinal (green arrowhead) neurons are visible in this projection. **(B)** Schematic of *NvLWamide-like::mcherry* neural subtypes are depicted as Tentacular (red), Pharyngeal (orange), Quadripolar (blue), Tripolar (magenta), Longitudinal (green), and Mesentery (purple). Oral end is to the left. Scale bar = 50μm.

**Figure 5.**
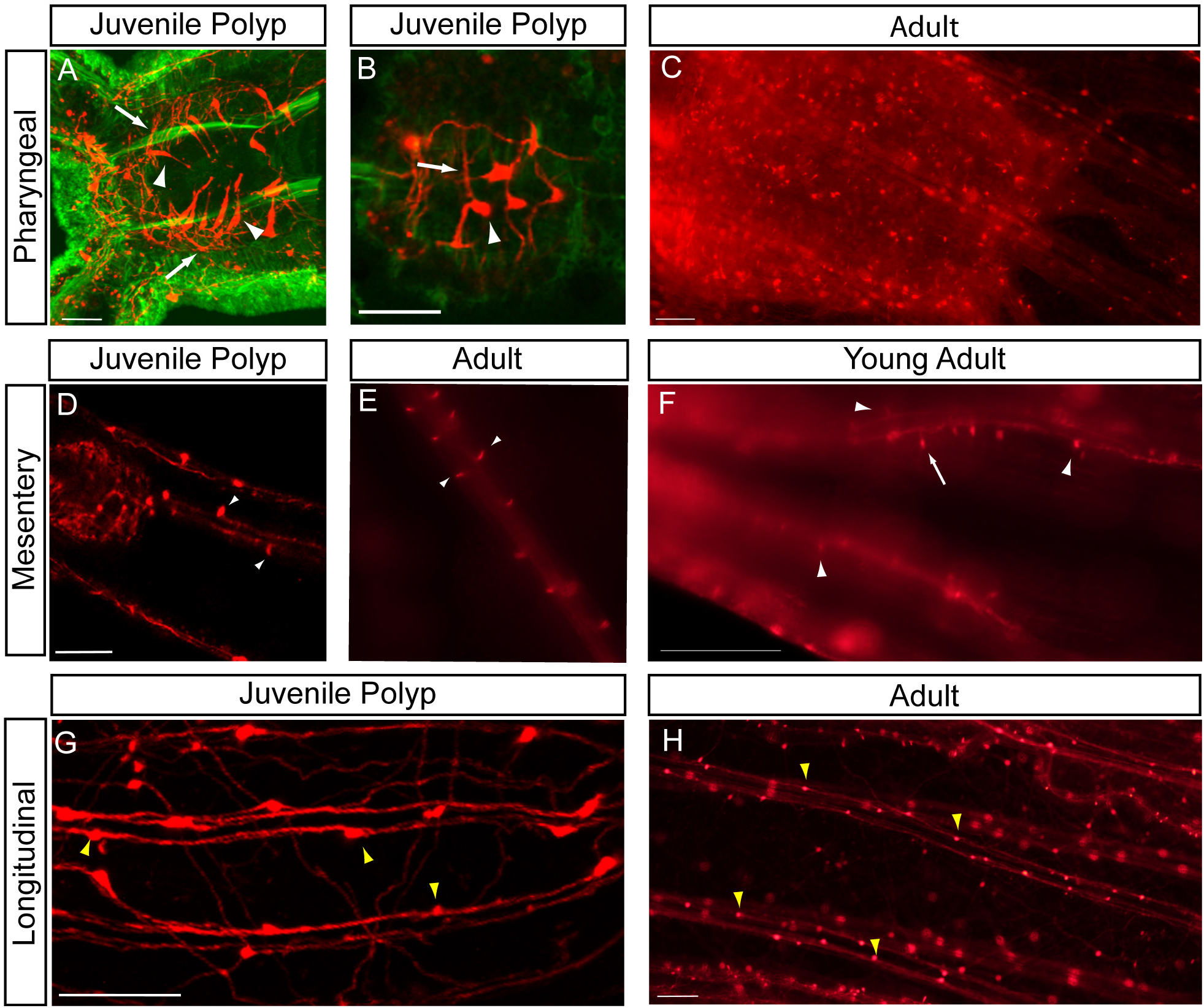
NvLWamide-like::mCherry transgenic line highlights multiple neuronal subtypes. **(A-C)** *NvLWamide-like::mcherry* expressing pharyngeal neurons (white arrows and arrowheads) are found in the juvenile polyp (A and B) and the adult (C) and wrap around the pharynx. **(D-F)** *NvLWamide-like::mcherry* positive neurons are observed in the mesenteries of both juvenile polyps (D) and adults (E and F), (D) Endodermal focal plane of a lateral view of a juvenile polyp shows mesentery cell bodies (arrowheads). (E) Endodermal view of fileted adult animal demonstrates that mesentery neurons are paired in adults. (F) Endodermal focal plane of a lateral view of an adult animal shows increased mesentery neurons and *NvLWamide::like* mesentery unipolar (arrow) and bipolar (arrowhead) neurons. **(G** and **H)** Longitudinal neurons (yellow arrowheads) are located in the endoderm and run in 2 parallel lines that follow the mesentery tracts in both juvenile polyps (G) and adults (H). The juvenile polyp shown in A and B was lightly fixed and co-stained with phalloidin (green). C-H represent live *in vivo* images of mCherry expression (red). In all images oral end is to the left. Scale bars = 20μm (A and B), 50μm (D and G) and 100μm (C, F, and H).

**Figure 6.**
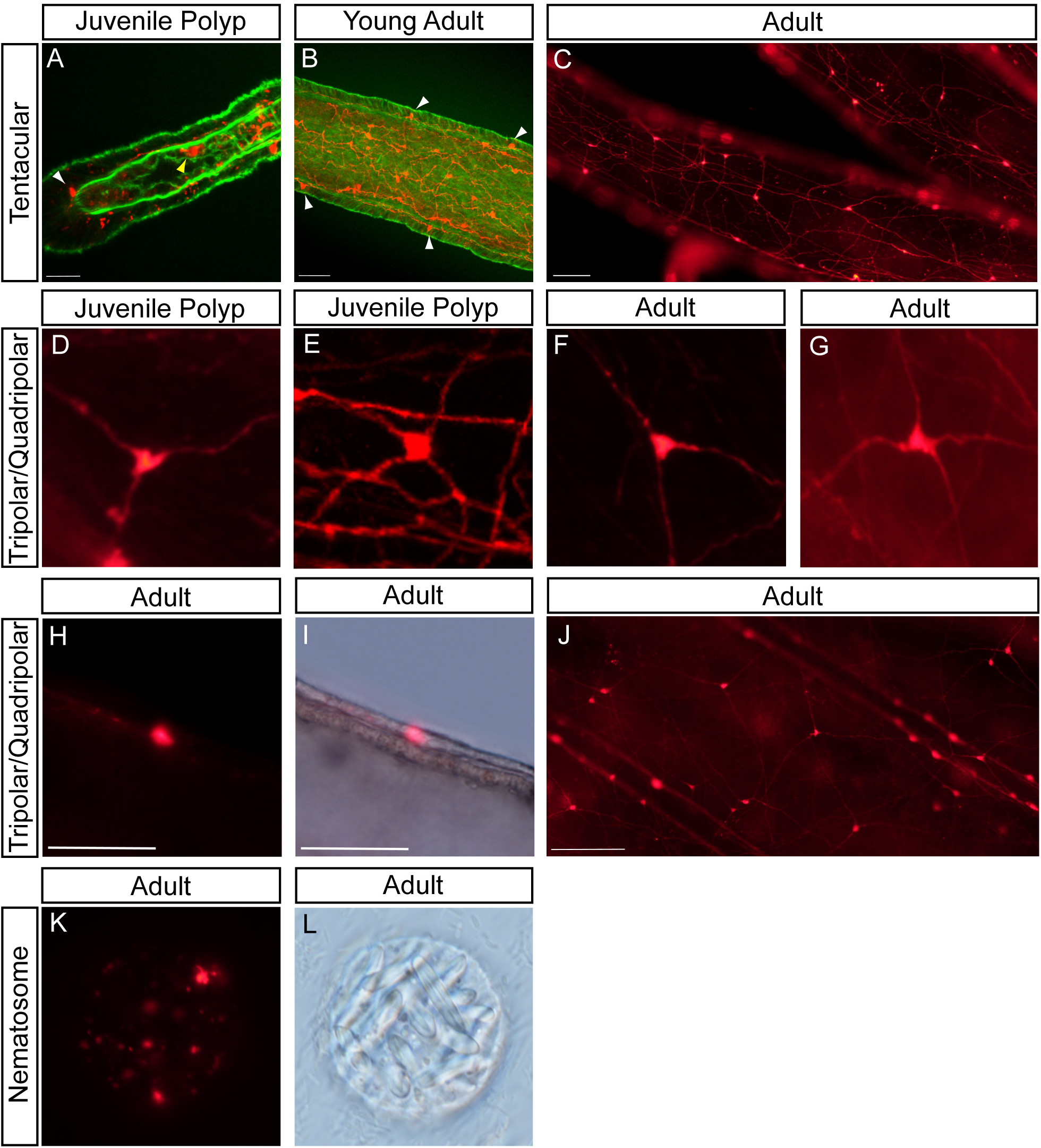
Ectodermal neuronal subtypes identified by *NvLWamide-like::mCherry* transgenic line. **(A-C)** Ectodermal *NvLWamide-like* tentacular neurons (white arrowheads) are observed in the basal ectoderm of the juvenile polyp (A), young adult (B) and adult (C). Neurons are located basally with projection that extend to other cell bodies (B and C) and appear to form a network of tripole neurons (C). *NvLWamide-like* tentacular neurons the have a sensory neuron morphology (B). Endodermal *NvLWamide-like* tentacular neurons are also found in the juvenile polyp (A, yellow arrowhead). **(D-J)** Neurons with a tripole and quadpole morphology are located throughout the entire ectodermal body column. Tripole neurons have 3 projections and are present in both the juvenile polyps (D) and adults (F, H-J). Quadpole neurons have 4 neurite projections and are present in juvenile polyps (E) and adults (G). Lateral view of an mCherry expressing tripole neuron (H). Same view and neuron as in H with DIC overlay, showing neuron is located in the ectoderm (I). By the adult stage a network of tripole and quadpole neurons is observed (J). **(K-L)** NvLWamide-like::mcherry expression is detected in the Nematosomes (K). DIC image of K showing the structure of the Nematosome (L). Samples in A and B were lightly fixed, and co-staining with phalloidin (green) highlights F actin filaments. C-L portray *in vivo* expression of mCherry captured in live animals. Scale bars = 20μm (A), 50μm (B, H-I) and 100μm (C and J).

### Pharyngeal neurons

*NvLWamide-like* pharyngeal neural somas are located within the pharyngeal ectoderm (Figure 4A, orange arrow; Figure 4B; Figure 5A, arrowheads). The neurites project out of the basal surface of the soma and encompass/wrap around the pharynx (Figure 5A and B, arrows). *NvLWamide-like* pharyngeal cell bodies span the pharyngeal ectoderm (Figure 5A, Supplemental Figure 1A, and Supplemental movie 1) and their apical surface appears to be exposed to the pharyngeal lumen. A view of the basal surface of the *NvLWamide-like* pharyngeal neurons suggests that initially the neurons are formed in a row (Figure 5B, arrowhead), each with 2 neurites that project orthogonally to the oral-aboral axis (Figure 5B, arrow). *NvLWamide-like* neurons contribute to the pharyngeal nerve mesh that surrounds the pharynx in the adult *Nematostella* (Figure 5C). By adult stages many *NvLWamide-like* pharyngeal neurons are detected throughout the pharynx and they do not appear to be restricted to a particular location. It is difficult to observe pharyngeal neurites in adult animals because the adult pharynx has robust red auto fluorescence. However, the *NvLWamide-like* pharyngeal neurites appear to maintain a mesh around the pharynx and do not appear to condense into a pharyngeal ring, which has been previously hypothesized to exist in *Nematostella* (Marlow et al., 2009). Based on the *NvLWamide-like* mRNA *in situ* and developmental time course in *NvLWamide-like::mcherry* transgenic animals (Figures 1 and 2) the pharyngeal neurons are born in the pharynx at early planula stages, coincident with or shortly after pharynx formation. As the animal matures and the pharynx increases size the number of pharyngeal neurons also grows.

### Mesentery neurons

*NvLWamide-like* mesentery neurons are aptly named for their location in the mesentery (Figure 4B; Figure 5D–F). Initially the juvenile polyp possesses few mesentery neurons (Figure 5D), but in feeding adults *NvLWamide-like* mesentery neurons are located along the entire length of the mesentery, (Figure 5E–F, arrowheads, and Supplemental Figure 1C). In adult *Nematostella* the soma seem to be in pairs (Figure 5E, arrowheads) mirrored on either side of the tract of neurites generated along the mesentery by these neurons (Figures 5E, 5F, and Supplemental Figure 1 C). We observe both unipolar (a single projection) (Figure 5F and Supplemental Figure 1 C, arrows) *NvLWamide-like* mesentery neurons and bipolar (two projections that project 180o opposite one another) (Figure 5F and Supplemental Figure 1 C, arrowheads) *NvLWamide-like* mesentery neurons. The projections emanate from the basal pole and fasciculate into bundles that project along the long axis of the mesentery (Figure 5F and Supplemental Figure 1 C). Because the tracts of neurites are immediately contacting the mesentery neural soma, it is difficult to estimate what percentage of the mesentery neurons are either uni-or bipolar. While we first observe *NvLWamide-like* mesentery neurons in the polyp stage, we speculate that their development occurs earlier, but that the delay in mCherry maturation makes identifying them in larval stages difficult.

### Longitudinal Neurons

The longitudinal *NvLWamide-like* neurons are endodermal and organized along the longitudinal neuronal tracts previously described to run above each of the eight mesenteries (Layden et al., 2016b; Nakanishi et al., 2012). The longitudinal tracts divide the body column of the polyp into 8 radial segments that extend the entire length of the oral aboral axis. Each longitudinal tract has two parallel bundles of fasciculated neurites. The *NvLWamide-like* longitudinal neural somas that reside within these tracts have bipolar projections that originate from the soma 180° to one another. (Figures 4, green arrowhead; Figure 5G and 5H, yellow arrowheads; Supplemental Figure 1B). The projections span the entire body length of the animal along the oral-aboral axis. The juvenile polyp usually has 2–3 *NvLWamide* positive cell bodies in each of the two parallel tracts present above each mesentery (Figure 5G), which translates to ~4–6 longitudinal neurons/radial segment (figure 7C). The mRNA *in situ* and developmental time course data show that longitudinal neurons are born in planula stages and neurites are already projecting prior to the tentacle bud stage (Figures 1; 2D). Longitudinal neurons show a dramatic increase in cell number in adult animals (Compare figures 5G to 5H and 7B to 7A). We quantified the number of longitudinal neurons in a number of animals (Figure 7C, light grey bars; Supplemental Table 1). Although the number is somewhat variable from animal to animal, the total number of neurons in each tract positively correlates with body length (Figure 7C), suggesting that the neuronal number is being controlled relative to the overall body length.

**Figure 7.**
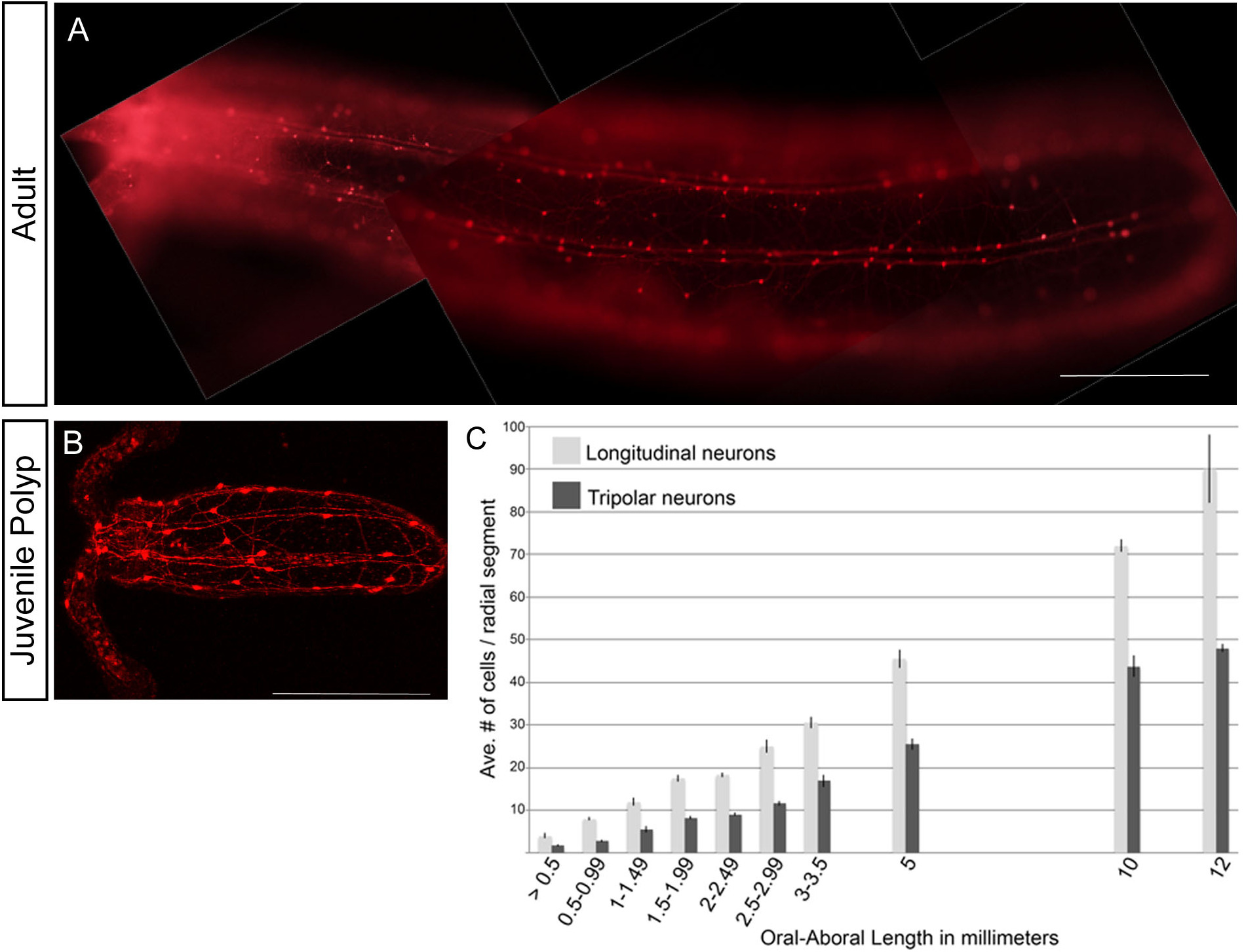
Comparison of *NvLWamide-like::mCherry* neurons present in juveniles and adults. **(A)** *NvLWamide-like* positive neurons in the juvenile polyp. **(B)** Adult *Nematostella* has a significant increase in the number of *NvLWamide-like::mCherry* positive neurons compared to the juvenile polyp, but no new neuronal *NvLWamide-like* positive subtypes are observed. In both animals oral end is to the left. Scale bars = 200μm. mCherry expression in A and B highlights *NvLWamide-like* positive neurons in live animals relaxed in 7.14% MgCL2 in 1/3X ASW. **(C)** Graph of the average number of longitudinal (light grey) and tripolar (dark grey) neurons present in each radial segment in animals of different lengths. Juvenile polyps are contained in the > 0.5 category. We were able to score two radial segments in each animal. N = 13, 5, 8,14,14, 9, 9, 4, 2,1 for > 0.5 to 12 mm respectively.

### Tentacular Neurons

*NvLWamide-like* tentacular neurons are first detected shortly after tentacle bud formation (Figure 4A, red arrow, and Figure 6A). Cell bodies are initially located both in the ectoderm (Figure 6A and B, white arrowhead) and endoderm (Figure 6A, yellow arrowhead) and are observed along the entirety of the proximal-distal axis of the tentacle. It is important to note that observing endodermal tentacular neurons is rare. A typical juvenile polyp tentacle has 1 to 3 cell bodies per tentacle. However, we have in one case observed seven cell bodies in one tentacle, suggesting that the exact number of neurons is somewhat variable. By adult stages the number of tentacular neurons dramatically increases and nearly all *NvLWamide-like* tentacular neurons are found in the ectoderm. It is unclear whether endodermal neurons migrate into the ectoderm or are lost by adult stages, but to date no trans-body layer migration has been described for *Nematostella* neurons. Tentacular neurites project in a fashion that appears to be similar to the tripolar neurons (described below) (Figure 6B–C).

### Tripolar and quadripolar neurons

The last two subtypes of neurons we identified are the *NvLWamide-like* tripolar and quadripolar neurons. These neurons are named for the number of projections from each soma. Tripolar and quadripolar neurons are found in the ectoderm. The soma for each neural type spans the entire depth of the ectoderm and their cell bodies are reminiscent of previously described sensory neurons (Figure 6D–J) (Marlow et al., 2009). The projections originate from the basal lateral surface and appear to project within or on the outer surface of the mesoglea. The juvenile polyp has at least 1 tripole neuron per radial segment, but we have observed up to 3 neurons per radial segment (Figure 4A, magenta arrowhead, and 6D; Figure 7C). The location of tripole neurons along the oral-aboral axis does not appear to be restricted to a particular region of the body column. At adult stages the number of tripole neurons increases, and similarly to the longitudinal neurons the increase in tripole neurons positively correlates with animal length (Figure 6F and J; Figure 7C).

At the early juvenile polyp stage we also observe a single *NvLWamide+* quadripolar neuron per animal (Figure 6E). Quadripolar neurons have four neurites that emanate from the soma and are similar to the tripolar in that they are located in the ectoderm. When detectable, quadripolar neurons can be first identified as early as 12 days post fertilization (dpf). Interestingly, unlike the tripolar neurons, these neurons appear to be more restricted to the oral portion of polyp trunk. While quadripolar neurons are also observed in the adult (Figure 6G) their numbers are not significantly increased, and their presence in the adult is difficult to detect. We cannot rule out that tripolar and quadripolar neurons are not the same neural subtype, but that there is some variability in the number of projections with three being significantly more common. We currently suspect that the tripolar and quadripolar neurons are the first neural cell types born at gastrula stages as that is the time we observe the most nascent ectodermal expression of *NvLWamide* (Figure 1). Interestingly, the number of *NvLWamide+* ectodermal cells decreases after gastrula stage suggesting that tripolar/quadripolar cells might be initially specified in excess and some of those cells undergo apoptosis. However, we cannot currently rule out that the early *NvLWamide-like* cells are unrelated to the tri and quadripolar cells, and that the tri and quadripolar cells are born later in larval stages.

### NvLWamide::mCherry is expressed in *Nematosomes*

Nematosomes are a *Nematostella* specific feature, comprised of a collection of both cnidocyte (stinging) cells and non-cnidocyte cells whose identity is unclear (Babonis et al., 2016). We detect NvLWamide::mCherry expression in the nematosomes along the entire length of the tentacle lumen (data not shown), in the nematosomes harvested from an egg mass (Figure 6K and L), and moving throughout the gut cavity of the polyp (not shown). Even though we did not detect nematosomes via mRNA *in situ,* previous reports detected *NvLWamide* expression in nematosomes by RNAseq analysis (Babonis et al., 2016).

### Summary of results

Our observations support stereotypy in the developing *Nematostella* nerve net. The juvenile polyp contains specific subtypes of *NvLWamide-like+* neurons, whose position, number, and neurite projection patterns show minimal variation from animal to animal. These data argue that although the nerve net appears unstructured when viewed in its entirety, it is comprised of neural subtypes that display significant organization when observed individually. Additionally, our data suggest that the neural cell types patterned during embryonic and larval development pioneer the neurite architecture. During adult growth the nervous system is modified in part by increasing numbers of individual subtypes in accordance with increasing body size.

## Discussion

### The *Nematostella* nervous system is organized and stereotyped at the subtype level

The *Nematostella* nervous system is often described as a “diffuse nerve net.” Here we examine *NvLWamide-like* expressing neurons, which represent a small subset of the nervous system to better visualize potential organization that might otherwise be masked if too many neurons are labeled. We observed reproducible neuronal subtypes: longitudinal, tripole, quadpole, pharyngeal, tentacular, and mesenetery. Moreover, we showed that the relative number of neurons for each neuronal subtype is similar from animal to animal upon completion of development, implying that neural number is patterned during development. These developmentally specified neural subtypes persist throughout the adult stage. However, the number of neurons in each subtype increases, and in the case of the longitudinal and ectodermal tripolar/quadripolar neurons there is a correlation between the numbers of neural bodies and body column length (Figure 7C). We hypothesize that many neuronal subtypes are stereotyped in terms of number of neurons and projections from those neurons. Together our data support the idea that the *Nematostella* nerve net is not a “structurally simple, diffuse net”, but instead specific patterns of neuronal subtypes and their projections are established during development and then neuronal number is modified in adult stages. Further characterization of the development and structure of other cnidarian developmental model nerve nets such as *Clytia hemisphaerica* and *Hydractinia* is necessary to confirm that this is a widespread phenomenon in cnidarian neurogenesis.

### Putative functions of subtypes expressing *NvLWamide-like*

We hypothesize that the tripolar, quadripolar, many of the tentacular and pharyngeal neurons are likely sensory neurons. They are all present in the ectoderm or ectodermally derived structures and have a cell body that spans the depth of the epithelial layer. The morphology of their soma is also consistent with previously predicted ectodermal sensory neurons (Marlow et al., 2009; Nakanishi et al., 2012). Mesentery and longitudinal neurons likely function as interneurons. Their projections fasciculate into either the mesentery or the previously described longitudinal neural tracts respectively (Marlow et al., 2009; Nakanishi et al., 2012). The longitudinal tracts contain a large number of neuronal projections, and are connected by commissural like structures (Supplemental Figure 3), which is consistent with a neuropil-like structure, where large number of interneuron like connections would be predicted to exist. Alternatively the positioning of the longitudinal and mesentery neurons could also reflect a role in contraction of myoepithelial cells as there is some evidence for LWamide family neuropeptides inducing muscle contractions in *Hydra* (Takahashi et al., 1997). We currently favor the hypothesis that the longitudinal neurons are interneurons because they are likely buried in the longitudinal tracts and not adjacent to the myoepithelial cells. However, we remain agnostic in regards to either hypothesis for the mesentery neurons.

### Development and patterning of the neuronal subtypes

The developmental mechanism(s) that pattern the *NvLWamide-like* subtypes remains unknown. Some evidence suggests that in *Nematostella* regional patterning likely interacts with neural programs to generate specific neural subtypes (Layden et al., 2012; Leclère et al., 2016; Marlow et al., 2013; Watanabe et al., 2014). This is most evident in the requirement of proper directive axis formation to pattern asymmetrically distributed GLWamide+ neurons (Watanabe et al., 2014), and in the expression of neural markers in the apical tuft, which requires proper establishment and maintenance of oral-aboral patterning programs (Leclère et al., 2016; Sinigaglia et al., 2015; 2013). Additional work has shown that regional boundaries with distinct gene expression profiles are established along the oral-aboral axis, which could be integrated into the generic neural patterning mechanisms previously identified to generate specific neural subtypes (Kusserow et al., 2005; Marlow et al., 2013). The tissue layer in which the neural cell resides and the temporal window in which the neural progenitor is born are additional factors that may contribute to neural subtype specification. To date our understanding of the temporal patterning in *Nematostella* is poor, but future work aimed at understanding how distinct neural fates arise should not ignore this component.

Our data also speak to the cellular mechanisms related to generating the appropriate number of each neural subtype from neural progenitor cells. Vertebrate neurogenesis occurs by overproducing neurons and neurons that fail to properly synapse with target cells do not survive. *Drosophila* ventral nerve cords form from hardwired neuroblasts that generate neural subtypes in an exact lineage so that there is essentially zero variability in regards to neural number between animals (Schmid et al., 1999). *Nematostella* neurogenesis appears to be in between these approaches. We observe stereotyped numbers of each neural subtype, but typically there is a 1–2 cell number variability for each cell type between animals and even between radial segments in the same animal. These data suggest that neural progenitor lineages are not hardwired. Additionally, with the exception of the reduction in ectodermal *NvLWamide-like* positive cells in the aboral ectoderm, it does not appear that there is a massive over production of *NvLWamide-like* neurons that then compete for a limited number of targets. We hypothesize the neural progenitor lineages are not hardwired in *Nematostella.*

*NvLWamide-like: :mcherry* expression describes a subset of the *Nematostella* nervous system. To fully understand the molecular and cellular programs that drive the development of diverse neural cell types, it will be necessary to generate new transgenic lines that identify additional subtypes of neurons in the *Nematostella* nervous system. The distinct neural subtypes will then need to be mapped to unequivocally establish their spatiotemporal origin during development, as well as the molecular program that specifies each cell type. Once we better understand the molecular programs controlling development of various neural cell types in cnidarians, we will then be poised ask questions regarding the relationship of bilaterian nervous systems to some or all of the cnidarian nerve net.

## Acknowledgements

We would like to acknowledge Uli Technau (University of Vienna, Austria) and Fabian Rentzsch (University of Bergen, Norway) for providing the pNvT vector necessary to generate the NvLWamide::mCherry transgene and the *Nvelav1::mOrange* transgenic strain. This research did not receive any specific grant from funding agencies in the public, commercial, or not-for-profit sectors.

